# BIN1 expression in the presynaptic compartment leads to isoform-specific synaptotoxicity

**DOI:** 10.1101/2025.08.11.669624

**Authors:** Erwan Lambert, Carla Gelle, Valentin Leclerc, Alejandra Freire-Regatillo, Nicolas Barois, Tommy Malfoi, Xavier Hermant, Florie Demiautte, Frank Lafont, Philippe Amouyel, Karine Blary, Sabine Kuenen, Chloé Najdek, Patrik Versteken, Dolores Siedlecki-Wullich, Pierre Yger, Jean-Charles Lambert, Devrim Kilinc, Pierre Dourlen

## Abstract

Alzheimer’s disease (AD) is characterized by a strong genetic predisposition and by an early loss of synaptic connectivity that strongly correlates with cognitive deficit. Some genetic determinants could contribute to synapse frailty toward AD pathology. However, the role of genetic determinants in AD pathogenesis remains poorly understood at the synaptic level. Here, we show that the expression of an isoform of the major AD susceptibility gene BIN1 in the presynaptic compartment results in synaptic loss. Using electrophysiology, we observed an early loss of synaptic transmission upon BIN1 isoform 1 (BIN1iso1) expression in *Drosophila* retinal photoreceptor neurons. This was not observed for the other human BIN1 isoforms tested, isoform 8 and isoform 9. Structural analysis of photoreceptor neuron synapses shows a strong accumulation of abnormally large vesicles in the presynaptic compartment, reminiscent of this same isoform-induced endosome defects in cell bodies. In addition, the expression of BIN1iso1 in motoneurons of the *Drosophila* neuromuscular junction alters the morphology of synaptic boutons, with a greater number and a smaller size of synaptic boutons, and the appearance of satellite boutons. As opposed to endosomal defects in cell body, modulating the Rab11 recycling endosome regulator did not prevent BIN1iso1 synaptotoxicity. To test if synaptic deficits are conserved in a mammalian model and to assert a presynaptic vs postsynaptic role for BIN1, we used rat primary neurons cultured in microfluidic devices that restrict gene expression modulation in particular neuron populations. We found a loss of synaptic connectivity only when expressing BIN1iso1 in the presynaptic compartment, which was confirmed by microelectrode array analysis. Together, our results suggest that BIN1 expression in the presynaptic terminal, but not the postsynaptic terminal leads to an isoform-specific, deleterious effect on synaptic integrity. BIN1 synaptotoxicity could contribute to the synapse loss observed early in AD. This supports the idea that genetic determinants could make synapses prone to failure in AD.

## Introduction

Alzheimer Disease (AD) is the most frequent neurodegenerative disease and cause of dementia worldwide. Besides the two pathognomonic protein aggregates, the extracellular senile plaques composed of amyloid-β peptide (Aβ) and the intracellular neurofibrillary tangles composed of phosphorylated tau, AD is characterized by progressive synaptic and neuronal losses. The synaptic loss strongly correlates with cognitive decline making synapse loss a central element of AD pathogenesis (DeKosky & Scheff, 1990; Mecca et al., 2022; Terry et al., 1991).

In addition, AD is considered to have a strong genetic component. Even in multifactorial cases, which represent the vast majority of AD forms, the heritability is estimated to reach 60-80% (Gatz et al., 2006). Recent advances in genetic studies have identified up to 76 genomic loci associated with the disease (Bellenguez et al., 2022). Genes responsible for the genetic signals in these loci are implicated somehow in the etiology of the disease. Therefore, some of them are likely involved in synapse loss. Accordingly, a genetically driven synaptic failure hypothesis has been raised for the disease (Dourlen et al., 2019). However, the potential synaptic roles of many AD risk genes are yet to be characterized. It is the case for *BIN1*, which is the susceptibility gene with the strongest association with AD after *APOE* (Bellenguez et al., 2022).

BIN1 has various functions in membrane trafficking, cytoskeleton regulation, DNA repair, cell cycle progression, and apoptosis (Dourlen et al., 2025; Prokic et al., 2014). *BIN1* is expressed in many organs, mainly in muscle and in the brain (Butler et al., 1997). In the brain, it is expressed mostly in oligodendrocytes, microglia and neurons (Adams et al., 2016; De Rossi et al., 2016; Marques-Coelho et al., 2021). *BIN1* has more than 10 isoforms. Some are ubiquituous, like BIN1 isoform 9 (BIN1iso9) and others are brain- or muscle-specific, like BIN1 isoform 1 (BIN1iso1) or BIN1 isoform 8 (BIN1iso8) respectively (Butler et al., 1997; Dourlen et al., 2025). In the brain, BIN1iso1 and BIN1iso9 are the most expressed (Crotti et al., 2019; Taga et al., 2020) and BIN1iso1 is considered neuron-specific (De Rossi et al., 2016).

BIN1 has been identified as a member of the Amphiphysin family of proteins and therefore involved in the regulation of endocytosis and the endolysosomal pathway (Butler et al., 1997; Ramjaun et al., 1997; Tsutsui et al., 1997; Wigge et al., 1997). Also named AMPH2, BIN1 shows sequence similarity with AMPH, a neuronal protein highly enriched in synaptic terminals and involved in synaptic vesicle endocytosis (David et al., 1996; Lichte et al., 1992; Shupliakov et al., 1997). BIN1 interacts with many components involved in clathrin-mediated endocytosis, namely dynamin, synaptojanin, AP-2, clathrin and endophilin (Drake & Traub, 2001; Leprince et al., 1997; McMahon et al., 1997; Micheva et al., 1997; Ramjaun et al., 1997). Furthermore, BIN1 has been visualized to be recruited to clathrin-coated pits in the sequence of events when the neck of the endocytic vesicle is formed just before vesicle scission (Taylor et al., 2012). BIN1 also contributes to fast endophilin-mediated endocytosis, a process regulating axonal growth *in vitro* in mouse hippocampal neurons (Ferreira et al., 2021). In Neuro2a cells, BIN1iso1 expression increases transferrin uptake (Ellis et al., 2012). However, in primary embryonic rat neurons, BIN1iso1 inhibits clathrin-mediated endocytosis resulting in smaller endosomes (Calafate et al., 2016). In human induced neurons, gain and loss of BIN1iso1 increases and decreases respectively the size of early endosomes (Lambert et al., 2022). This effect of BIN1iso1 overexpression is conserved when expressed in *Drosophila* neurons and leads to neurodegeneration, which can be prevented upon inhibition of the early endosome regulator Rab5 and upon activation of the recycling endosome regulator Rab11 (Lambert et al., 2022). Of note, regulation of the endosome size is of interest for AD as endosomal enlargement is one of the first AD pathological hallmark (Cataldo et al., 1997, 2000).

Intracellular trafficking is crucial for synapses; in the presynaptic compartment it ensures the correct generation and release of synaptic vesicles and in the postsynaptic compartment it is required for the correct recycling of neurotransmitter receptors. From *Drosophila* to mammals, BIN1 is localized at synapses (Leventis et al., 2001; Ramjaun et al., 1997; Zelhof et al., 2001). Whether it is localized more in the presynaptic compartment or in the postsynaptic compartment depends on the experimental model and study (De Rossi et al., 2020; Schürmann et al., 2020). At the functional level, in the postsynaptic compartment, *BIN1* has been shown to regulate the recycling of AMPA receptors at the plasma membrane *in vitro* (Schürmann et al., 2020). In the presynaptic compartment, loss of *BIN1* has been shown to regulate presynaptic vesicular release in hippocampal excitatory synapses *in vivo* (De Rossi et al., 2020). Therefore, the role of BIN1 at the synapse remains poorly understood and it is essential to address this question if we aim at better understanding its role in AD pathogenesis.

For this purpose, we investigated the synaptic role and potential toxicity of human BIN1iso1, BIN1iso8 and BIN1iso9. We used the highly tractable *in vivo* model *Drosophila*, in which human BIN1 functions are conserved. This is exemplified by human BIN1 being able to rescue the locomotor deficits of endogenous *Drosophila* BIN1 mutants (called Amphiphysin) (Lambert et al., 2022). We first analyzed the synaptotoxicity of human BIN1 isoforms expression in the retina photoreceptor neurons. We confirmed our results in third instar larval glutamatergic neuromuscular junction (NMJ), a synaptic reference model in *Drosophila* (Menon et al., 2013). We further tested the toxicity of BIN1iso1 and BIN1iso9 in rat primary neuronal cultures using microfluidic devices and multielectrode (MEA) arrays, which enabled us to assess pre- and postsynaptic roles of BIN1 in synaptic structure and transmission in the mammalian context.

## Material and Methods

### Drosophila genetics

Flies were raised on a standard medium (Nutri-fly BF, Genesee Scientific, San Diego, CA, USA) at 25°C under a 12h/12h day/night cycle. The UAS-BIN1iso1, UAS-BIN1iso9, UAS-BIN1iso8, UAS-dAmphA and rh1-Gal4 lines were described previously (Dourlen et al., 2012; Lambert et al., 2022). Other stocks were obtained from the Bloomington *Drosophila* Stock Center (BDSC, Bloomington, IN, USA): UAS-Luciferase (#35788), UAS-mCD8::GFP (#27400), UAS-GFP (#35786), UAS-Rab11::GFP (#8506), Nsyb-Gal4 (#42714).

### Western blotting

*Drosophila* heads (10 heads per condition) were dissected and crushed in 30µL of NuPAGE LDS buffer (NP0008, NuPAGE, Novex, Life Technologies) supplemented with reducing agent (NP0009, NuPAGE, Novex, Life Technologies). Samples were centrifuged (8500g, 10min, 4°C) and the supernatants were stored at -80°C. Once thawed, they were denatured by heating for 10 minutes at 85°C, then plated and separated on a 4-12% bis-tris acrylamide gel (NuPAGE, Novex, Life Technologies) in 1X MOPS buffer (NP0001-02, NuPAGE, Novex, Life Technologies). Samples were transferred to nitrocellulose membrane (NuPAGE, Novex, Life Technologies) using the Biorad Trans-Blot transfer system kit (Biorad) according to the supplier’s technical recommendations (7 min, 2.5A, 25V). The quality of migration and transfer was checked by staining with Ponceau Red 0.2% (Sigma).

Membranes were incubated for 1h at room temperature in 1X TNT (Tris 0.15M, Nacl 1.5M, Tween20 0.5%) supplemented with 5% skimmed milk powder to block aspecific sites, and labeled with primary antibody in SuperBlock Tween 20 blocking buffer (37563, Thermo Scientific) overnight at 4°C. This incubation was followed by three 10-minute rinses with 1X TNT to remove aspecifically bound antibodies. The membrane was then incubated with the secondary antibody for 2 hours, followed by 3 further 10-minute rinses with 1X TNT to remove excess of secondary antibodies. The antibodies used were: anti-BIN1 primary antibodies (BIN1 99D, 05-449, Merck Millipore, RRID:AB_309738, 1/2500, BIN1 ab27796, abcam, RRID:AB_725699, 1/1000), anti-α-tubulin primary antibody (α-tubulin DM1A, T9026, Sigma, RRID: AB_477593, 1/5000), anti-actin primary antibody (Sigma-Aldrich Cat# A2066, RRID:AB_476693), anti-GFP primary antibody (anti-GFP, G1544, Sigma, RRID:AB_439690, 1/4000) and HRP-conjugated secondary antibodies (Jackson ImmunoResearch, anti-mouse 115-035-003, RRID:AB_10015289 and anti-rabbit 111-035-003, RRID:AB_2313567, 1/10000). Immunoreactivity was revealed by incubation with ECL (WBLUC0500, Immobilon Classico Western HRP Substrate, Millipore) and the chemiluminescence imaged with the Amersham Imager 600 camera (GE lifesciences, GE Healthcare, USA). Relative quantification was performed using Fiji software.

### Electroretinography

Electroretinograms (ERGs) were recorded in flies immobilized on a glass microscope slide using liquid Pritt glue. For the recordings, glass pipettes (borosilicate, 1.5 mm outer diameter; Hilgenberg) were filled with 3M NaCl and placed in the thorax, as a reference, and over the fly’s eye, lightly penetrating the cornea for the recordings. Responses to a repetitive light stimulus (1 s) given by a green light-emitting diode were recorded using AxoScope 10.5 software and analyzed using Clampfit 10.5 software (Molecular Devices). Recordings were amplified using a Warner DP311 AC/DC amplifier (Warner Instruments) and digitized using MiniDigi 1A (Molecular Devices). Raw data traces were analyzed with Igor Pro 6.36 (Wavemetrics).

### Electron microscopy

Electron microscopy was performed as described previously (Lambert et al., 2022). After dissection, *Drosophila* eyes were fixed in 1% glutaraldehyde, 4% paraformaldehyde, 0.1M sodium cacodylate buffer (pH 6.8) 30 min at room temperature and then overnight at 4°C. After washing, eyes were post-fixed in 1% OsO4 and 1.5% potassium ferricyanide for 1h, then with 1% uranyl acetate for 45 min, both in distilled water at room temperature in the dark. After washing, they were dehydrated with successive ethanol solutions. Eyes were infiltrated with epoxy resin (EMbed 812 from EMS) and were mounted in resin into silicone embedding molds. Polymerization was performed at 60°C for 2 days. Ultrathin sections of 70-80 nm thickness were observed on formvar-coated grid with a Hitachi H7500 TEM (Milexia, France), and images were acquired with a 1 Mpixel digital camera from AMT (Milexia, France).

### NMJ dissection and immunostaining

Third instar larvae were placed on a Petri dish coated with sylgard, a bicomponent silicone (Sylgard^TM^ 184 Elastomer Base, Dow Corning), in a drop of cold Schneider’s medium (Schneider’s *Drosophila* Medium (1X), 21720-024, Gibco), to prevent the larvae from drying out and to stun them, making the work easier. The larvae were then pinned between the posterior spiracles and at the level of the pharyngeal apparatus, dorsal side up. Using dissecting scissors, we cut the dorsal cuticle along the anteroposterior axis in-between the two dorsal tracheal trunks to open larvae. We performed two additional horizontal incisions at the level of anterior and posterior spiracles. The muscular walls were deployed on either side of the vertical section to reveal the interior. We then removed the internal organs. Once the dissection was complete, we replaced the Schneider medium with 100 µL of 4% PFA fixation solution (Electron Microscopy Sciences, 15712) in 1x PBS (70011-036, Gibco) per larva for 20 minutes. We then performed three 10-minute washes with PBT (PBS 1x with 0.1% Triton-X100, AppliChem A4975). Saturation was carried out with 5% NGS (Normal Goat Serum, Jackson ImmunoResearch, 005-000-121) in PBT for 30 minutes. Primary antibodies were deposited overnight at 4°C in PBT 0.1% NGS 5%. The following antibodies were used: anti-HRP488 (123-545-021, Jackson ImmunoResearch, RRID:AB_2338965, 1/400), anti-Disc Large (Dlg 4F3, DSHB, RRID:AB_ 528203, 1/250), anti-Bruchpilot (nc82, DSHB, RRID:AB_2314866, 1/250) and anti-BIN1 (99D, Merck Millipore, RRID:AB_309738, 1/250). The following day, we performed three 10-minute washes with 0.1% PBT. Following this, we incubate the larvae with secondary antibodies in PBT 0.1% NGS 5%, 2 hours at room temperature: phalloidin (anti-F-actin Alexa 555, A34055, ThermoFisher Scientific), Alexa 633 Goat anti-mouse (A21052, Life Technology, RRID:AB_ 2535719) and Dye Light 405 Donkey anti-rabbit (711-475-152, Jackson ImmunoResearch, RRID:AB_ 2340616). We then performed three 10-minute washes with 0.1% PBT followed by a rinse with 90% glycerol in 1X PBS. NMJs were imaged with a Gataca Systems disk spinning microscope (Gataca Systems, France) equipped with a Yokogawa CSU-W1 head (Yokogawa, Japan), a Prime95B camera (Photometrics, Teledyne Technologies Inc., USA) and a Ti-2 inverted stand (Nikon, Japan). To quantitatively analyze NMJs, we manually delineated each synaptic bouton (including satellite boutons if present) one by one using FIJI (NIH, Bethesda, MD) and measured their area. These values were statistically analyzed using R software.

### Microfluidic device preparation

Microfluidics masters were fabricated at the Institute of Electronics, Microelectronics and Nanotechnology, Lille, France using two-step photolithography as previously described (Blasiak et al., 2015). For synaptic connectivity analysis, we used a three-chamber design where the side chambers house neuronal cell bodies, and the central (or synaptic) chamber houses neurites arriving from the side chambers and the synaptic connections they form (Kilinc et al., 2020). The device consists of a 450-µm-wide central channel flanked by two 750-µm-wide side channels. All three channels are *ca.* 100 µm high. The left side channel (termed presynaptic) and the central channel (termed synaptic) are interconnected via 4-mm-high, 450-mm-long parallel microchannels that narrow from an entry width of 10 µm to an exit width of 3 µm. The right-side channel (termed postsynaptic) and the synaptic chamber are also interconnected via parallel microchannels with identical dimensions, except that they were 75-µm-long. 4-mm-high polydimethysiloxane pads were replica molded. Access wells were punched at the termini of the synaptic chamber and of the side channels using 3-mm and 4-mm biopsy punches (Harris Unicore), respectively. The devices were permanently bonded to 24 × 50 mm glass coverslips (Menzel) via O2 plasma (Diener, Ebhausen, Germany). Prior to cell culture, the devices were sterilized under ultraviolet light (Light Progress, Anghiari, Italy) for 20 min, treated with 0.1 mg/mL poly-D-lysine (Sigma) for at least 2 h, followed by 20 μg/mL laminin (Sigma) treatment for 2 h. The devices were then washed once with PBS prior to neuron seeding.

### Primary neuron culture

Culture media and supplements were purchased from Thermo Fisher unless specified otherwise. Primary neurons were obtained from P0 (post-natal day 0) rats according to established procedures (Sartori et al., 2019). The hippocampi were isolated from the cortex and washed with ice cold dissection medium (Hanks’ balanced salt solution supplemented with HEPES, 1% sodium pyruvate, and 0.5% penicillin/streptomycin) and treated with 2.5% Trypsin at 37°C for 10 min. The trypsin was then inactivated using an MEM/FBS medium (Minimum Essential Media supplemented with 10% heat inactivated fetal bovine serum, 1% GlutaMAX, 3% D-glucose (Sigma), 0.8% Minimum Essential Media vitamins and 0.5% Pen/Strep) and incubated with 5 mg/mL DNase (Sigma) for 1 min followed by 3 washes with MEM/FBS medium. The cells were then dissociated in culture medium (Neurobasal A, supplemented with 1% GlutaMAX and 2% B27 neural supplement with antioxidants) before being centrifuged at 200 ×*g* for 8 min. Cells were then resuspended in culture medium, counted and plated at 100,000 cells/cm^2^ density in the pre- and post-synaptic chambers. The next day, the medium in reservoirs was replaced with fresh culture medium. The 0.1% ethylenediaminetetraacetic acid (in H2O) was added to the Petri dishes containing microfluidic devices to minimize evaporation. The cultures were maintained in a tissue culture incubator (Panasonic; Osaka, Japan) at 37°C and 5% CO2 for up to 21 days.

### BIN1 isoform overexpression

Overexpression constructs were obtained from Gene Art (Thermo Fisher) based on pLenti6.3/Ubc/V5-DEST A244 backbone vectors with a CMV promoter (Life Technologies, Carlsbad, CA): BIN1iso1 (NM_009668), BIN1 isoform 9 (NM_139349), and an overexpression control vector (Mock). On day 7 *in vitro* (DIV7), neurons in the pre-synaptic or post-synaptic chambers were transduced as previously described (Eysert et al., 2021). To avoid the transduction of neurons in the opposite chamber, a hydrostatic pressure gradient is formed across the microchannels separating the synaptic chamber and the target cell chamber. Lentiviral particles were diluted in pre-warmed culture medium containing 2 µg/mL Polybrene (hexadimethrine bromide; Sigma). Media from all wells were collected in a common tube. A total of 20, 15 and 25 µL of the collected medium were added to the reservoirs of the target cell chamber, synaptic chamber and opposite cell chamber, respectively. Cells were transduced at a multiplicity of infection (MOI) of 2 by adding 10 µL of virus suspension to one of the wells of the target cell chamber. Neurons were incubated with viral particles for 6 h before the wells were topped-up with the remaining collected medium. In a separate experiment, cells in the pre- and postsynaptic chambers were transduced with lentiviral vectors encoding LifeAct-Ruby (pLenti.PGK.LifeAct-Ruby.W; RRID:Addgene_51009) and LifeAct-GFP (pLenti.PGK.LifeAct-GFP.W; RRID:Addgene_51010), respectively, to demonstrate that viral transductions were restricted to the intended cell chamber.

### Immunocytochemistry of primary neurons

To quantify synaptic connectivity, we fixed the neurons on DIV14 in 4% paraformaldehyde (PFA) for 15 min, washed 3× with PBS, and permeabilized for 5 min with 0.3% Triton X-100. Cells were incubated with 5% normal donkey serum for 2 h at RT before overnight incubation with the following primary antibodies: BIN1 (Millipore Cat# 05-449, RRID: AB_309738), Homer1 (Synaptic Systems Cat# 160 004, RRID: AB_10549720), MAP2 (Synaptic Systems Cat# 188 006, RRID: AB_2619881). The cells were then washed 3× with PBS and incubated with the following secondary antibodies raised in donkey: AlexaFluor-conjugated AffiniPure Fragment 405, 594, or 647 (Jackson ImmunoResearch). The cells were then washed 3× with PBS and incubated with a Cy2-tagged monoclonal antibody against Synaptophysin 1 (Synaptic Systems Cat# 101 011C2, RRID: AB_10890165).

### Microscopy and synaptic connectivity analysis

Microfluidic devices were imaged using a Zeiss LSM880 confocal microscope, equipped with a 63× 1.4-NA objective. Images were analyzed with Imaris software (Bitplane, Zürich, Switzerland) by reconstructing Synaptophysin I and Homer puncta in 3D. The volume and position information of all puncta were processed using a custom Matlab (MathWorks, Natick, MA) program. This program assigns each postsynaptic spot to the nearest presynaptic spot (within a distance threshold of 1 μm) and calculates the number of such assignments for all presynaptic puncta. The percentage of presynaptic spots assigned by at least one postsynaptic spot was systematically used as a readout for synaptic connectivity (Kilinc et al 2020).

### Microfluidic device integration to MEAs

We used 256-electrode MEA chips (256MEA100/30iR-ITO-w/o, MultiChannel Systems, Reutlingen, Germany) where electrodes form a 16×16 Cartesian grid, except for the corners where the tips of large, triangular reference electrodes are positioned. As previously done for 60-electrode MEA chips (Lefebvre et al., 2024), we adapted the microfluidic device layout by adding a triangular chamber between the flow channels connecting synaptic and postsynaptic chambers to their respective medium reservoirs. This chamber matches the tip of one reference electrode such that the reference electrode is fluidically connected to the rest of the device. To minimize the effect of this reference electrode chamber on fluid flow in the aforementioned channels, we used sets of parallel microchannels (18 µm width; 7 µm separation) for the connections. MEA-adapted microfluidic devices were prepared in the same way as the regular microfluidic devices and align-bonded to MEA chips using O2 plasma.

To position similar numbers of microelectrodes underneath pre- and postsynaptic chambers, we use a mechanical tool to precisely align the microfluidic device over the MEA chip (Lefebvre et al., 2024). Briefly, after O2 plasma exposure, the MEA chip was placed on a manual stage allowing rotation and two-axis translation in the *xy*-plane (Newport; Irvine, CA). The microfluidic device was held above the MEA chip with a vacuum pen (Virtual Industries Inc.; Colorado Springs, CO) that was attached to manual *z*-axis translational stage. Using binoculars, the microfluidic device was then aligned with the MEA chip in a way that pre- and postsynaptic chamber were covered by *ca.* 50 electrodes each. The device was then lowered onto the MEA chip until contact was established, and the integrated device was kept at 70°C for 2 min to strengthen the bond. The MEA-integrated microfluidic devices were placed in 100 mm polystyrene Petri dishes and sterilized and coated in the same way as the regular microfluidic devices. A 35 mm Petri dish containing sterile milliQ water was placed next to the MEA chips to minimize evaporation.

### MEA recordings

At DIV19, the spontaneous neuronal activity in each MEA chip was recorded using a MEA2100-System equipped with a single 256-channel headstage and an integrated temperature controller (MultiChannel Systems, Reutlingen, Germany). First, the headstage was preheated to 37°C. A plastic ring with ethylene-propylene membrane was placed over the microfluidic device to ensure sterility during transport and recordings. The MEA chip was brought on the headstage and left there for 5 min for the neurons to adjust to the new environment. Spontaneous activity was recorded from all channels for 5 min at 25 kHz sampling rate. The recordings were exported to .h5 files for further analysis. For each MEA-integrated microfluidic device, a chip map was generated that lists the electrodes corresponding to pre- and post-synaptic chambers.

### MEA analysis

#### Spike Sorting

The extracellular recordings were analyzed using SpikeInterface (Buccino et al., 2020), and the SpyKING CIRCUS 2 spike sorting pipelines, conceived as an extension of our spike sorting toolbox (Yger et al., 2018). Briefly, the spike sorting procedure to extract the single unit activities starts with a preprocessing step, as follows: bandpass filter (frequency [150Hz, 7kHz]), then a common median reference filter (to remove shared fluctuations across channels) before whitening (to remove shared noise across channels). Then, negative threshold crossings are detected, aligned, and a subset is projected into a lower dimensional space via Principal Components (see (Yger et al., 2018) for details). Density-based clustering is then launched on these projected spikes to detect the templates, *i.e.*, the motifs regularly occurring in the data. Once the dictionary of templates is constructed, a proper orthogonal matching pursuit algorithm (Tropp & Gilbert, 2007) reconstructs the signal as a linear sum of these templates to identify all the spike times. After spike sorting, only the putative units with a refractory period violation less than 0.1 and a signal-to-noise ratio more than 2 were kept as valid neurons.

#### Position of the units

The putative positions of the units kept after spike sorting are estimated from the extracellular templates, assuming that units could be considered as monopoles (Scopin et al., 2024; Varol et al., 2021). All the units that would be found within the presynaptic chamber are labeled as belonging to the *pre* population, while those found within the postsynaptic chamber are labeled as *post*. Note that while all electrodes were analyzed simultaneously during the spike sorting process (to make use of spatiotemporal information to disambiguate sources), very few units were found between the two chambers, in accordance with the experimental setup. We analyzed the ratio of *post* units to *pre* units to identify potentially abnormal devices. To this end, we used Matlab (Mathworks, Natick, MA) to generate a probability plot for normal distribution, using detected unit numbers for all devices (Supplementary figure 5). Based on this plot, devices with |*z*-score| > 1.5 were considered outside the normally distributed post:pre unit number ratio and discarded from further analysis. Note that this threshold corresponds to 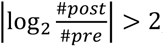, *i.e.*, a maximum of 4-fold difference in unit numbers between the two chambers was allowed.

#### Drive between *pre* and *post* populations

To assess the directionality of the functional connectivity between pairs of units, we defined a metric called the *pre-post-drive* that measures to what extent the units in the *pre* population are leading the activity of the units in the *post* population. The assumption behind such a metric is that assuming monosynaptic, direct connections between *pre* and *post* neurons, one must see an increase in activity in the *post* population following spikes emitted by the *pre* population. To do so, for each pair of unit *i* belonging to the *pre* population and of unit *j* belonging to the *post* population, we measured the cross-correlogram *CC*_*_i_pre__*,*_j_post__*_ (*t*) of their spiking activity with a given time bin *T*_*bin*_ = 2 *ms* and during a time window *T*_*window*_ = 20 *ms*. The *drive* Φ(*i_pre_*, *j_post_*) was then computed as the ratio of the integral of *CC*_*_i_pre__*,*_j_post__*_ (*t*) for positive times (*i.e.*, how much the activity of *i_pre_* drives the one of *j_post_*) divided by the integral of *CC*_*_i_pre__*,*_j_post__*_ (*t*) for negative times (*i.e.*, how much the activity of *j_post_* drives the one of *i_pre_*). If the two units were firing independently, as Poisson sources, then Φ(*i_pre_*, *j_post_*) should be close to 1. If, on the other hand, Φ(*i_pre_*, *j_post_*) is larger than 1, it means that, on average, spikes of *i_pre_* is more likely to precede the ones in *j_post_*. Such an asymmetry in the cross-correlogram is often used to infer putative directional connectivity (Aertsen & Gerstein, 1985). Then, at the population level, the *drive* between *pre* and *post* populations is simply obtained as Φ(*pre*, *post*) = *mean*_*i*∈*pre*,*j*∈*post*_Φ(*i_pre_*, *i*_*post*_). Data points beyond median ± 3 median absolute deviations (MAD) were deemed as outliers and excluded from subsequent statistical analysis.

### Statistical analysis

Statistical information is available in the figure legends. Two-tailed statistical tests were used. Hypothesis testing was performed with the Kruskal Wallis test followed by a Mann-Whitney comparison, or by a post-hoc Dunn multiple comparison test (Holm method), or with Tukey-Kramer corrections. Statistical analyses were performed using Matlab (Mathworks, Natick, MA), R 3.6.0 (R Core Team (2019). R: A language and environment for statistical computing, R Foundation for Statistical Computing, Vienna, Austria. URL https://www.R-project.org/), especially with the FSA package (Ogle et al., 2023), and RStudio 1.2.1335. For box plots, the middle segment, lower and upper hinges represent the median, first quartile and third quartile respectively. The upper whisker extends from the upper hinge to the largest value within 1.5 x IQR of the hinge (where IQR is the interquartile range, or the distance between the first and third quartiles). The lower whisker extends from the lower hinge to the smallest value located at a maximum of 1.5 * IQR from the hinge. Differences with a *p*-value <0.05 were considered significant.

## Results

### BIN1iso1 altered synaptic transmission of adult photoreceptor neurons in *Drosophila*

To assess the effect of BIN1 isoforms on synaptic transmission, we investigated the electrophysiological response of *Drosophila* photoreceptor neurons expressing BIN1 isoforms upon light illumination. Flies expressed BIN1iso1, BIN1iso8, BIN1iso9, dAmphisoA (the longest endogenous BIN1 isoform) and Luciferase (used as a control) in the outer photoreceptor neurons at day2, day8-9 and day15-16 after birth. The expression was driven by a Rh1 promoter, active just before birth, after photoreceptor neuron development, and by the Gal4/UAS system (Brand & Perrimon, 1993). We checked BIN1 isoforms and dAmphisoA protein expression by western blot using two antibodies. The UAS constructs are inserted in the same genomic location and therefore exhibit similar expression (Lambert et al., 2022). All the constructs were expressed as expected (Supplementary figure 1). The ab27796 antibody cross-reacted partially with dAmphisoA whereas the 99D antibody did not detect dAmphisoA and more strongly detected BIN1iso1. On these flies, we performed electroretinograms (ERGs). Upon a 1 sec-long orange light stimulation, retinas usually exhibit a depolarization and two transient currents, called the ON transient and the OFF transient, at the initiation and termination of the light stimulation respectively (Figure 1A). The ON and OFF transients reflect the synaptic transmission activity of the photoreceptor neurons at the level of the lamina, the outermost part of the optic lobe. On day 2, we did not observe any alteration of the electrophysiological activity whatever the BIN1 isoforms. This indicated that all photoreceptor neurons were normal at birth. On day 8-9, we observed a significant reduction of the amplitude of the ON and OFF transients specifically for flies expressing BIN1iso1 (Figure 1A, B). On day 15-16, the amplitude of all electrophysiological parameters, the ON and OFF transients and the depolarization, were strongly reduced for retinas expressing BIN1iso1 (Figure 1A, B). It was not observed for retinas expressing BIN1iso8, BINiso9 and dAmphisoA. The amplitude of the OFF transient was even slightly increased in these conditions. The loss of all electrophysiological parameters on day 15-16 for BIN1iso1 likely reflected neurodegeneration and loss of the photoreceptor neurons that have been shown in these flies (Lambert et al., 2022). The loss of the ON and OFF transients on day 8-9, observed here, showed in addition that BIN1iso1 expression altered synaptic transmission before neurodegeneration.

**Figure 1:**
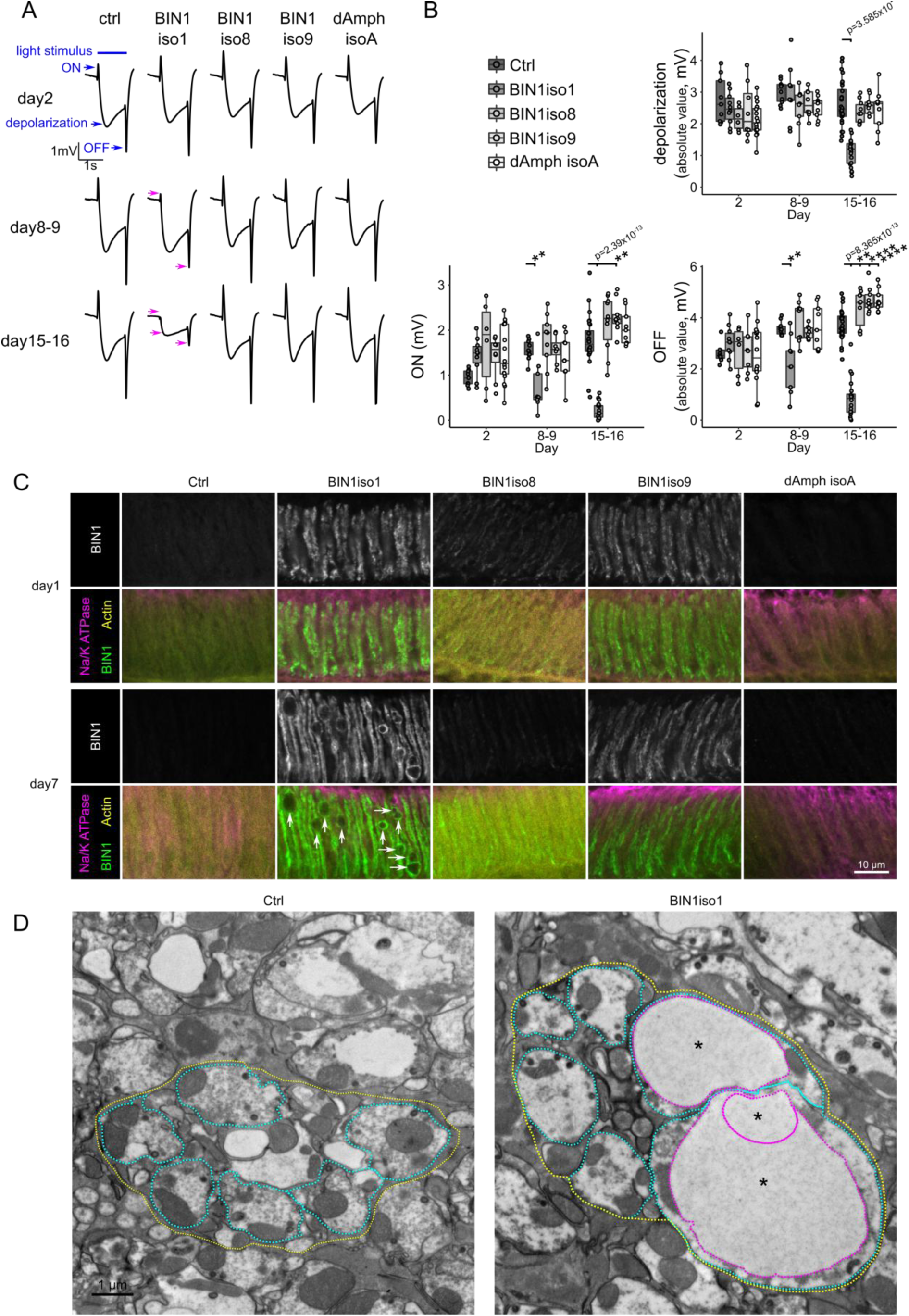
BIN1iso1 disrupted synaptic transmission of *Drosophila* photoreceptor neurons with age. **A.** Electroretinograms of *Drosophila* expressing BIN1iso1, BIN1iso8, BIN1iso9, dAmphisoA and Luciferase (control) under a rh1 driver at day 2, day 8-9 and day 15-16. Specifically, for BIN1iso1, we observed a loss of ON and OFF transients (magenta arrows) at day 8-9 followed by a loss of all electrophysiological parameters, (depolarization, ON and OFF transients, magenta arrows) at day 15-16. **B.** Quantification of the depolarization and ON and OFF transients (Kruskal-Wallis ANOVA followed by Wilcoxon rank-sum test). **C.** Immunofluorescence of the synaptic extremities of photoreceptor neurons expressing BIN1iso1, BIN1iso8, BIN1iso9, dAmphisoA and Luciferase (as a control) under a rh1 driver at day 1 and day 7. BIN1 staining shows the localization of BIN1iso1, BIN1iso9 and BIN1iso8 at the presynaptic terminal at day 1 and day 7. The neuron-specific plasma membrane Na/K ATPase and the actin staining are used to label these structures. At day 7, presynaptic extremities of BIN1iso1-expressing photoreceptor neurons accumulate big vesicles (arrows). **D.** Electron microscopy analysis of synaptic terminals of photoreceptor neurons expressing Luciferase and BIN1iso1 at day 15. Six synaptic terminals (cyan dotted lines) are gathered into a lamina cartridge (yellow dotted lines). Expression of BIN1iso1 resulted in the presence of giant vesicles (*, magenta dotted lines).

To understand the isoform-specificity of the result, we wondered whether all BIN1 isoforms were localized at the photoreceptor neuron axon synaptic terminals in the lamina. By immunofluorescence in 1-day-old whole-mount *Drosophila* retina, we observed that BIN1iso1, BIN1iso9 and to a lesser extent BIN1iso8 were all localized to the synaptic terminals (Figure 1C). This remained true in 7-day-old flies. Therefore, BIN1iso1 synaptotoxicity did not originate from an aberrant localization of this isoform at the synaptic terminals. In addition, we observed in 7-day-old flies a strong accumulation of vesicles in the presynaptic terminals for the BIN1iso1 condition. Some of these vesicles were very big, even bigger than the size of synaptic terminals (Figure 1C, arrows). We further analyzed the synaptic terminal structure using electron microscopy (Figure 1D). We confirmed the presence of abnormal giant vesicles. These vesicles had a single membrane and did not have any specific content. We could see mitochondria and capitate projections as in the control. Finally, we observed some degenerating synaptic terminals (Supplementary figure 2). Overall, the vesicles, observed in the synaptic terminals, perfectly corresponded to the ones exhibiting endosomal markers, described in photoreceptor neuron cell bodies (Lambert et al., 2022). Therefore, they likely originate from a defect in intracellular trafficking and are responsible for the synaptic transmission defects and synaptic terminal degeneration.

### BIN1iso1 altered *Drosophila* larval neuromuscular junction

We wanted to further assess the role of BIN1 isoforms on synapse integrity and tested the effect of their expression in third instar larval type Ib neuromuscular junctions (NMJ). It is a large glutamatergic synapse between a motor neuron and a muscle cell. Expression of BIN1 isoforms and dAmphisoA was performed thanks to a Nsyb driver. We checked the expression of the protein by western blot in adult head originating from the larvae, as the Nsyb driver remains active in adulthood, and observed similar results as in the eye with the rh1 driver (Supplementary figure 3). For each larva, we observed the same NMJ, located on muscle cells 6/7 in abdominal segment A2. We labelled the actin cytoskeleton (Phalloidin), the neuron-specific horseradish peroxidase (HRP) epitope and Disc large (Dlg), the orthologue of PSD95, to visualize muscles, motor neuron axon terminals and postsynaptic compartments respectively (Figure 2A).

**Figure 2:**
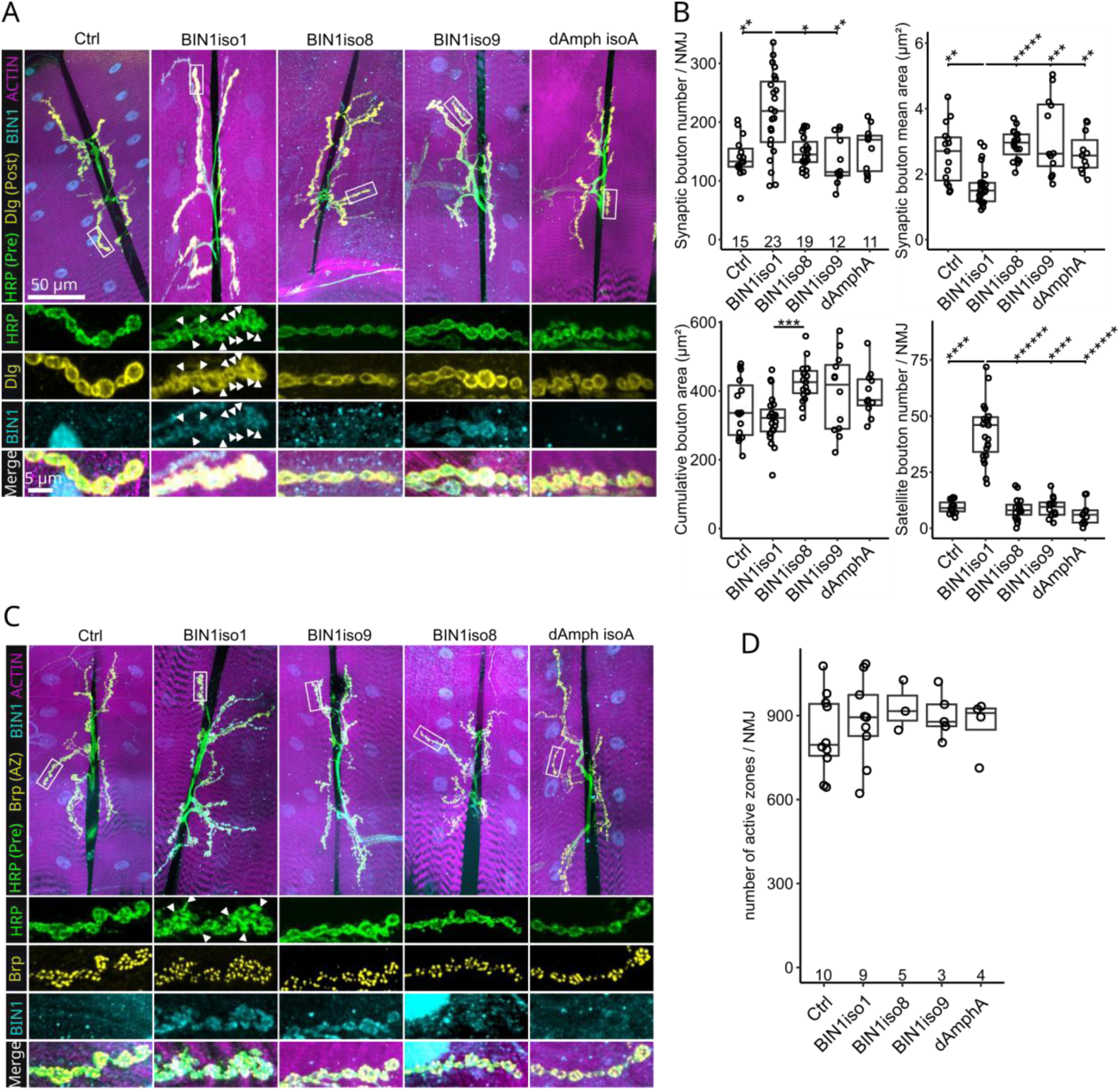
BIN1iso1 caused morphological alterations in *Drosophila* NMJs without affecting the number of active zones. **A.** Immunofluorescence of NMJs from 3^rd^ instar larvae expressing BIN1iso1, BIN1iso8, BIN1iso9 and dAmphisoA and Luciferase (control) in motoneurons with the Nsyb promoter. HRP, Dlg and actin labeling are used to label motor neuron terminals, postsynaptic compartments and muscle cells, respectively. Arrowheads show satellite boutons. **B.** Quantification of NMJ morphometric parameters: the number of boutons per NMJ, the mean area of single bouton per NMJ, the cumulative area of all boutons per NMJ and the number of satellite boutons per NMJ. The number of NMJ analyzed per condition are indicated at the bottom of the top left graph. Data were analyzed with a non-parametric Kruskal Wallis test followed by post-hoc Dunn’s multiple comparison test (Holm method, * p<0.05, ** p<0.01, *** p<1.10^-3^, **** p<1.10^-4^, ***** p<1.10^-5^, ****** p<1.10^-6^). **C.** Immunofluorescence of NMJs from 3^rd^ instar larvae expressing BIN1iso1, BIN1iso8, BIN1iso9 and dAmphA and Luciferase (control) in motoneurons with the Nsyb promoter. HRP, Brp and actin labeling are used to label the motor neuron terminals, active zones and muscle cells, respectively. Arrowheads show satellite boutons. **D.** Quantification of the number of active zones. Numbers at the bottom of the graph indicate the number of quantified NMJs. Non-parametric Kruskal Wallis analysis did not show any statistically significant difference between conditions.

In larvae expressing luciferase (used as a control), BIN1iso8, BIN1iso9 and dAmphisoA, the innervation pattern appeared to be normal, with large, well-rounded, well-contoured Ib-type synaptic boutons and a stereotyped branching pattern (Figure 2A). It indicates that these isoforms did not impact NMJ morphology. In contrast, BIN1iso1-expressing synaptic boutons looked shredded, clustered in places, and their contours were not as sharp and defined as the control boutons. When quantifying the boutons, it translated in a significantly higher number (∼225 boutons per NMJ versus ∼150 for the control) of smaller boutons (∼1.75 µm² versus ∼2.5 µm² for the control) in the BIN1iso1-expressing condition (Figure 2B). In addition, in this condition, we were able to see a significantly higher number of satellite boutons (∼40 satellite boutons per NMJ compared with ∼10 for the control), *i.e.*, boutons budding off from a bouton present in the axis of the branch, or boutons budding off from neuronal connections between two boutons. We did not observe any significant change when summing all individual bouton areas (cumulative area), nor when quantifying the number of active zones using a bruchpilot (brp) staining (Figure 2C, D). Active zones are synaptic vesicle-rich compartments in the presynaptic boutons, where precisely orchestrated molecular interactions control the fusion of synaptic vesicles with the presynaptic membrane. It indicates that despite the strong NMJ morphological alterations, the total size of the NMJ remained constant between conditions. Thanks to a BIN1 staining, we could detect BIN1iso1, BIN1iso9 and to a lesser extent BIN1iso8 in the motor neuron terminals and synaptic boutons (Figure 2A, C), indicating that the specificity of BIN1iso1 synaptotoxicity does not result from a different addressing of BIN1 isoforms in motor neurons.

### Rab11 expression did not prevent BIN1iso1-induced morphological changes of the *Drosophila* larval neuromuscular junction

It is known in the PR neuron model, that BIN1iso1 leads to degeneration via a defect in early endosome trafficking, and that the activation of recycling endosomes via Rab11 alleviates the neurodegeneration (Lambert et al., 2022). Because in this work, BIN1iso1-associated synaptotoxicity exhibited endosomal-like vesicles and because the intracellular trafficking is crucial for synaptic function, we hypothesized that BIN1iso1 synaptotoxicity could also originate from an intracellular trafficking defect. To test this hypothesis, we expressed the recycling endosome regulator Rab11 and assessed if it was possible to restore the BIN1iso1-induced phenotype of NMJs to a wild-type appearance.

Wild-type NMJs exhibited normal morphology with round, well-defined synaptic boutons (Figure 3A). Expression of BIN1iso1 resulted again in NMJs featuring a higher number of smaller synaptic boutons, with a higher number of satellite boutons (Figure 3A, B). Overexpression of Rab11 alone did not affect the morphology of synaptic neurons but significantly decreased the number of synaptic boutons (Figure 3A, B). In this background, overexpression of BIN1iso1 still increased the number of synaptic and satellite boutons, although without changing the mean area of single boutons, and the increase in synaptic bouton number was significantly lower than in the control background (Figure 3A, B). This indicated that Rab11 overexpression did not clearly prevent the effect of BIN1iso1 on NMJs and suggested more an additive effect rather than an epistatic one between the overexpression of BIN1iso1 and Rab11. We checked that the staining of BIN1was similar between the BIN1iso1 alone and the BIN1iso1+Rab11 conditions, which we confirmed by western blot (Supplementary figure 4). Overall, these results suggested that BIN1iso1 synaptotoxicity is rather independent of Rab11-controlled endosomal trafficking.

**Figure 3:**
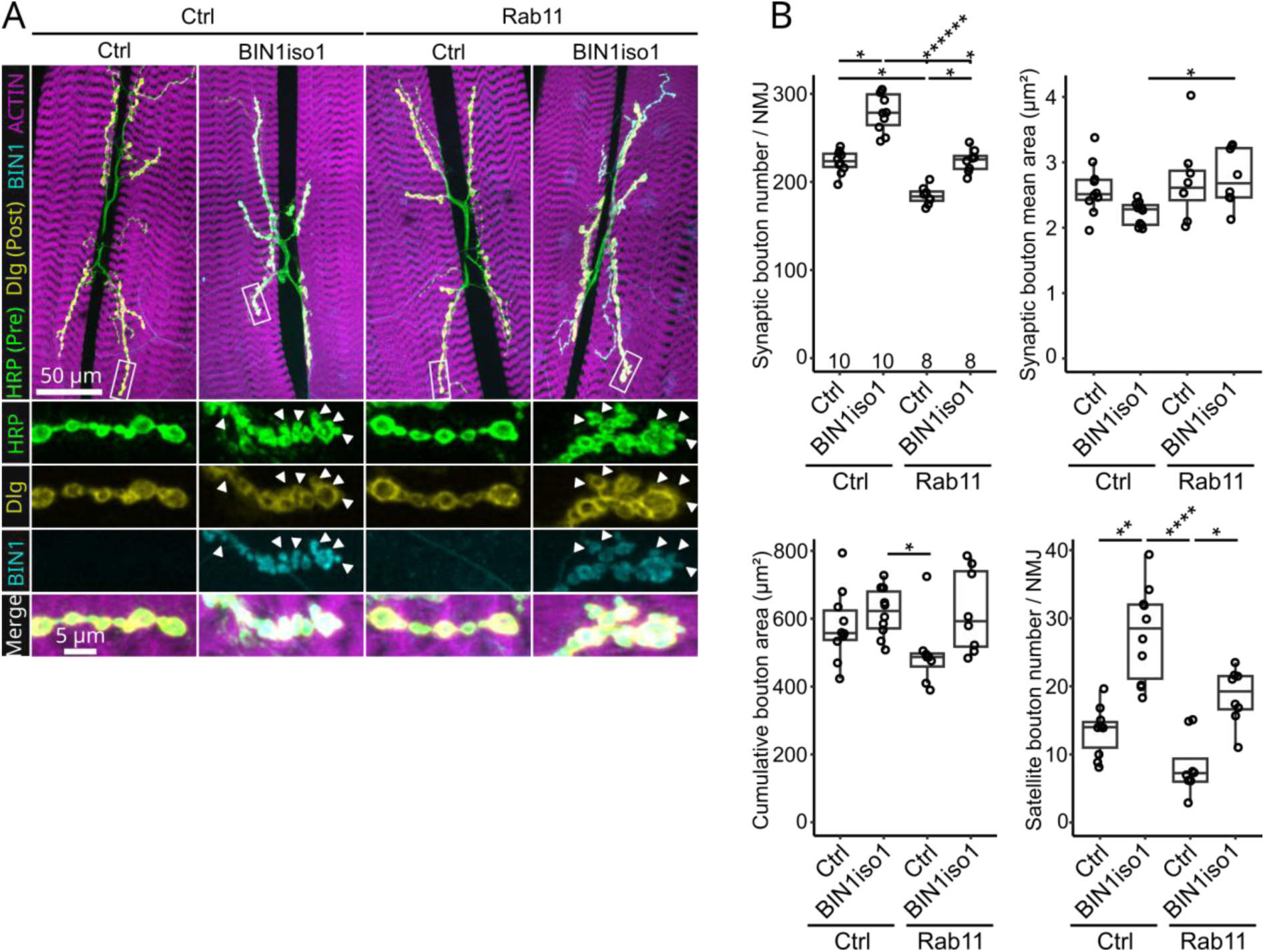
Rab11 did not prevent BIN1iso1-induced morphological defects at the larval NMJs. **A.** Immunofluorescence of NMJ from 3rd instar larvae expressing BIN1iso1 and/or Rab11 in motoneurons (Nsyb driver). mCD8::GFP and luciferase are used as controls respectively for Rab11::GFP and BIN1iso1. HRP, Dlg and actin labeling are used to label motor neuron terminals, post-synaptic compartments and muscle cells respectively. **B.** Quantification of NMJ morphometric parameters: the number of boutons per NMJ, the mean area of single bouton per NMJ, the cumulative area of all boutons per NMJ and the number of satellite boutons per NMJ. The number of NMJ analyzed per condition are indicated at the bottom of the top left graph. Data were analyzed with a non-parametric Kruskal Wallis test followed by post-hoc Dunn multiple comparison test (Holm method, * p < 0.05, ** p < 0.01, **** p<1.10^-4^, ****** p<1.10^-6^).

### BIN1iso1 is toxic for rodent synapses only when expressed in the presynaptic compartment

We next sought to determine if the isoform-specific effect of BIN1iso1 on synapses could be replicated in a mammalian system. To this end, we cultured rat hippocampal neurons in custom-design, three-chamber microfluidic devices that fluidically isolate synapses from their cell bodies and permit modulation of gene expression exclusively in pre- or postsynaptic chambers (Kilinc et al., 2020). By employing two sets of parallel microchannels of different length to connect presynaptic and postsynaptic chambers to the (central) synaptic chamber (Figure 4A), these devices ensure that the synaptic chamber receives dendrites only from the postsynaptic chamber and contains 83.2±6.1% of all synaptic connections between pre- and postsynaptic chambers (Kilinc et al., 2020). We plated primary neuron cultures in pre- and postsynaptic chambers and transduced them on DIV7 with lentiviruses encoding for human BIN1iso1, BIN1iso9 or a scrambled control vector (Mock), either in the presynaptic or in the postsynaptic chamber. At DIV14, we fixed the cultures and immunostained them against BIN1, somatodendritic marker MAP2, presynaptic marker Synaptophysin and postsynaptic marker Homer (Figure 4B). We applied our distance-based synaptic connectivity analysis workflow, which assigns each postsynaptic puncta to the nearest presynaptic puncta within a distance threshold (Eysert et al., 2021; Kilinc et al., 2020; Saha et al., 2024). Overexpression of BIN1iso1 –but not BIN1iso9– in the presynaptic chamber –but not in the postsynaptic chamber– induced synaptotoxicity in mature hippocampal neurons as evidenced by a decrease in the fraction of presynaptic spots assigned by at least one postsynaptic spot (Figure 4C). This corroborates our findings in *Drosophila* NMJ that BIN1 induces isoform-specific perturbations at the presynaptic compartment.

**Figure 4:**
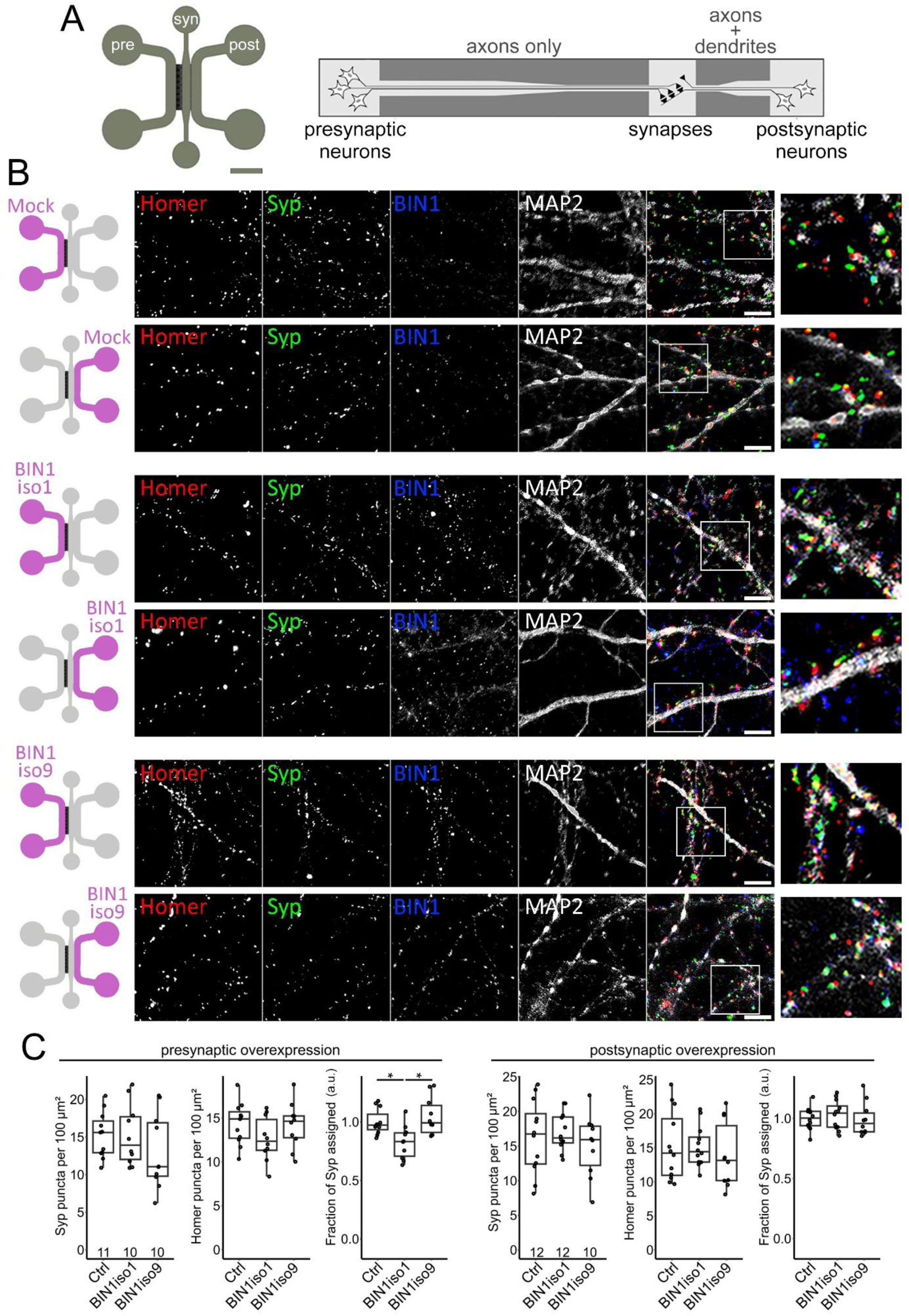
Presynaptic overexpression of BIN1iso1 induces synaptotoxicity in hippocampal neurons. **A.** Scheme of the custom-design, three-chamber microfluidic devices used to culture primary rat hippocampal neurons. Scale bar = 2 mm. **B.** Representative images of the synaptic network at DIV14. Overexpression of Mock (control), BIN1iso1 or BIN1iso9 vectors in the pre- or postsynaptic neuron chamber is indicated in the left schemes (scale bars = 5 µm). **C.** Quantification of Synaptophysin (Syp) and Homer spot densities and the fraction of Syp spots assigned by at least one Homer spot, as a measure for synaptic connectivity between the two neuronal populations. In box plots, data points correspond to device means. Fraction of Syp assigned was normalized by the mean of the control group. N = 3 independent cultures each for presynaptic and postsynaptic expression. Kruskal–Wallis ANOVA with Tukey-Kramer correction. * *p* < 0.05. N/S: not significant. a.u.: arbitrary units.

### Presynaptic BIN1iso1 overexpression decreases network-level functional connectivity

Analysis of synaptic connectivity by immunofluorescence provides inherently limited information on the functionality of the neuronal network. We therefore assessed network-level synaptic connectivity in three-chamber microfluidic devices integrated to MEA chips (Figure 5A-C) – based on the experimental model we recently developed in the context of Aβ-induced synaptotoxicity (Lefebvre et al., 2024). We analyzed the spiking activity via an established spike sorting algorithm (Yger et al., 2018), which identifies putative units based on their spatiotemporal firing patterns and calculated the cross-correlation for each unit pair that can be formed between units in the pre- and postsynaptic chambers (Figure 5D-F). As a measure for functional connectivity, we defined a *pre-post-drive* parameter by taking the ratio of right-hand-side integral to left-hand-side integral of the cross-correlation histogram for each such unit pair (Figure 5G) (see Materials and Methods for details).

**Figure 5:**
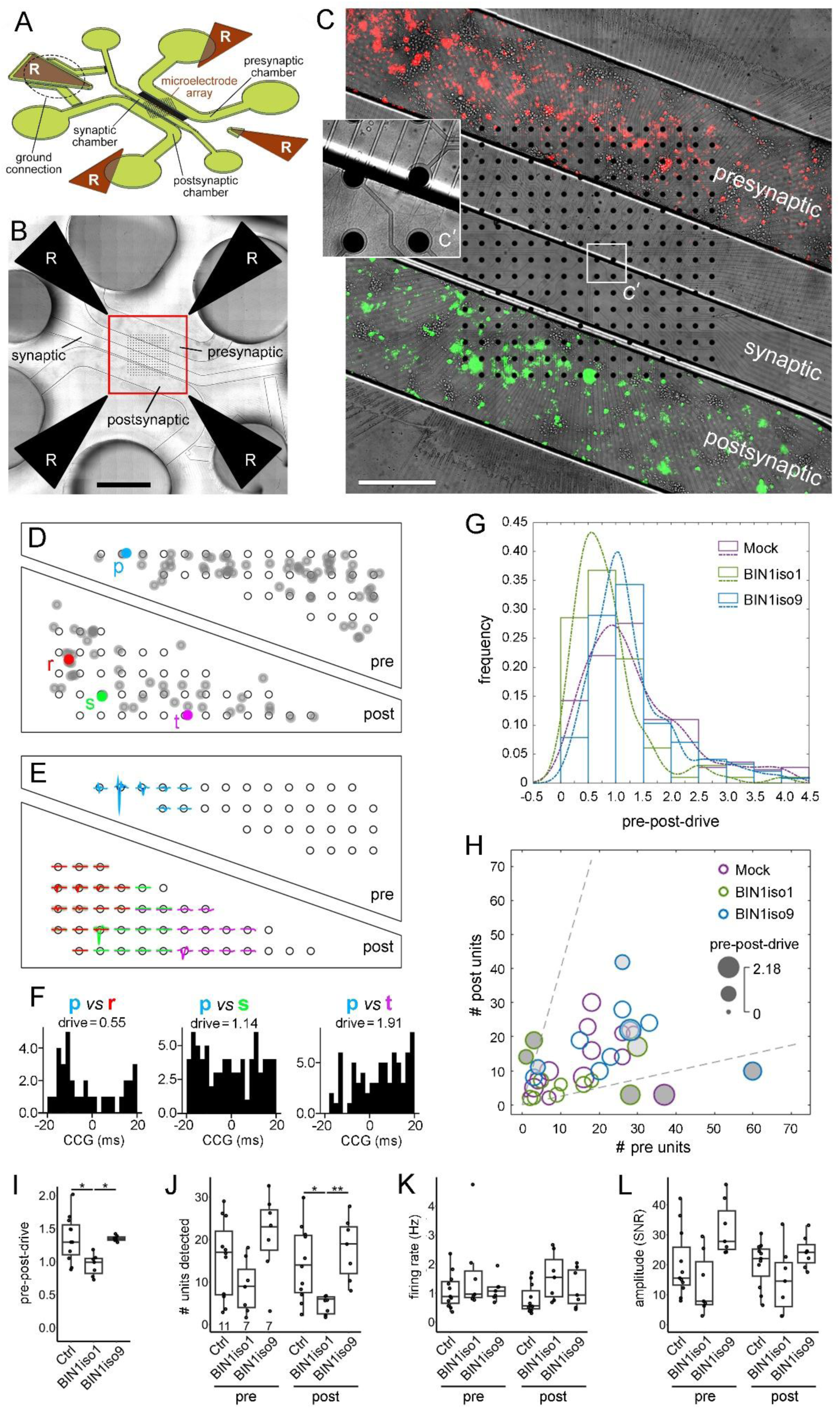
Presynaptic overexpression of BIN1iso1 perturbs the network-level connectivity in hippocampal neurons. **A.** Schematics of the three-chamber microfluidic device integrated with a 256-electrode MEA. Reference electrodes are marked with “R”. **B.** Mosaic image of the MEA-integrated microfluidic device housing rat hippocampal neurons. Cells in the pre- and postsynaptic chambers were lentivirally transduced to express LifeAct-Ruby (red) and PSD95-YFP (green) vectors, respectively, to demonstrate isolated transductions in cell chambers. Scale bar = 2 mm. **C.** Boxed area in panel B at higher magnification. Scale bar = 500 µm. Zoomed area (C’) is 3.4× magnified. **D.** A representative map of units identified through the spike-sorting algorithm (gray). One presynaptic unit and three postsynaptic units are color-coded for demonstration purposes. **E.** Spike waveform patterns corresponding to these four color-coded units. **F.** Cross-correlograms and the calculated *pre-post-drive* parameters for the three pre-post unit pairs that can be formed using these four units. **G.** Pre-post-drive histograms for representative microfluidic devices (one device per condition). Lines show the corresponding kernel distribution fits. **H.** Bubble chart showing *pre-post-drive* as a function of numbers of units detected in the pre- and postsynaptic chambers. The unit number ratios were normally distributed among all devices (see Supplementary figure 5), except for those with |*z*-score|>1.5 (dark gray). This threshold corresponds to a 4-fold difference in the unit number ratio (broken lines). Outlier data points according to median±3×MAD are included (light gray). **I-L.** *Pre-post-drive* (I), number of detected units (J), mean firing rate for these units (K) and mean amplitude for these units (L) (L) as a function of BIN1 isoform (or Mock vector) expression in the presynaptic chamber. In box plots, data points refer to individual devices. N = 6 independent cultures. Kruskal–Wallis ANOVA with Tukey-Kramer correction. * *p* < 0.05; ** p < 0.005. N/S: not significant.

We made a total of 40 recordings (14 Mock, 15 BIN1iso1, 11 BIN1iso9, expressed only in the presynaptic chamber) using primary neuronal cultures from 7 independent cultures. For 5 devices (1 Mock and 4 BIN1iso1) the algorithm was not able to identify any units in the presynaptic chamber (that passed the defined quality criteria). In addition, we discarded a total of 5 recordings (1 Mock, 3 BIN1iso1, 1 BIN1iso9) as they fell outside the normal distribution of post:pre unit number ratio (Supplementary figure 5). For the remaining recordings (12 Mock, 8 BIN1iso1, 10 BIN1iso9), we first assessed whether the number of units detected by the spike sorting algorithm correlated with *pre-post-drive*. When all conditions were pooled, the pre-post-drive parameter did not correlate with the number of units detected in the presynaptic chamber (R^2^ = 0.136), number of units detected in the postsynaptic chamber (R^2^ = 0.078) or with the total number of units detected (R^2^ = 0.122) (Supplementary figure 6). This finding suggests that the *pre-post-drive* parameter can be used as a read-out for network-level connectivity, independently of the number of units detected in the microfluidic device.

We then compared the experimental groups in terms of detected unit numbers and pre-post-drive. To this end, we first plotted pre-post-drive as a function of numbers of detected units in each chamber (Figure 5H). For clarity, we included in this plot the outlier data points as defined by median ± 3×MAD pre-post-drive (1 Mock, 1 BIN1iso1, 3 BIN1iso9). Interestingly, the pre-post-drive was significantly lower in the BIN1iso1 group compared to the Mock and BIN1iso9 groups (Figure 5I). In fact, for the BIN1iso1 condition, the pre-post-drive parameter was very close to 1, indicating that the presynaptic chamber was no longer driving the postsynaptic chamber, in contrast with the other conditions. The number of units in the postsynaptic chamber was also lower for the BIN1iso1 group when compared to Mock and BIN1iso9 groups despite the fact that the overexpression was restricted to the presynaptic chamber (Figure 5J). This suggests that the postsynaptic activity is (in part) driven by the pre-to-post synaptic connections, which are perturbed by the expression of BIN1iso1 in the presynaptic chamber. The number of units in the presynaptic chamber was also decreased, but this is not reflected in Figure 5J due to the exclusion of 4 out of 15 devices with no presynaptic units, as this precludes cross-correlation analysis. Finally, despite these detrimental functional changes caused by BIN1iso1, neither the mean firing rate (Figure 5K) nor amplitude (signal-to-noise ratio) (Figure 5L) were affected. In summary, the MEA results based on the spontaneous activity of live neurons recapitulate our microscopy-based findings and confirm that presynaptic BIN1iso1 overexpression leads to a loss of network-level synaptic connectivity.

## Discussion

Our findings demonstrate that the overexpression of BIN1iso1 induces synaptic dysfunction both morphologically and functionally across invertebrate and vertebrate models. In *Drosophila* photoreceptor neurons, BIN1iso1 impaired synaptic transmission prior to the complete loss of electrophysiological response, suggesting that synaptic dysfunction is an early and possibly primary event in BIN1-mediated toxicity. Notably, this effect was specific to isoform 1, as neither BIN1iso9 nor BIN1iso8 recapitulated the phenotype. Morphologically, BIN1iso1 expression resulted in an altered NMJ architecture characterized by an increased number of synaptic boutons, many of which were smaller in size and appeared as satellite boutons. This structural remodeling was accompanied in the photoreceptor synaptic terminals by an accumulation of endosome-like vesicles, pointing to a disruption of vesicular trafficking. These vesicles were reminiscent of the ones seen in cell bodies of degenerating photoreceptors, which could be rescued by gain-of-function of Rab11, a key regulator of recycling endosomes. In this work, we found that overexpression of Rab11 only partially rescued in an additive manner the synaptic defects caused by BIN1iso1, suggesting a slightly different mechanism at the synaptic level. We then extended our observations to mammalian models using rat hippocampal neurons. BIN1iso1 overexpression in the presynaptic compartment significantly reduced synaptic density, while postsynaptic expression had no detectable effect. This presynaptic specificity was again isoform-dependent, as BIN1iso9 expression did not affect synaptic density. Functionally, presynaptic BIN1iso1 expression led to disrupted synaptic transmission and altered network activity, further highlighting a conserved and isoform-specific synaptotoxic effect.

We showed in this work an early synapse toxicity for BIN1iso1. Because synapses are known to be affected early in AD and their loss correlates with cognitive decline, BIN1, a risk gene for the disease, may contribute to this process (DeKosky & Scheff, 1990; Mecca et al., 2022; Terry et al., 1991). This is supported by some variants of BIN1, associated with AD and shown to increase the expression of BIN1iso1 (Chapuis et al., 2013). Other evidence suggests that BIN1 expression decreases in the later stages of AD (Marques-Coelho et al., 2021; Saha et al., 2024). This could be due to the death of high-expressing neurons, leaving only low-expressing neurons. Alternatively, the decline in BIN1 expression may be causal in disease progression. In this context, therapeutic strategies aimed at increasing BIN1 activity need to be carefully controlled to avoid exacerbating neurotoxicity due to BIN1iso1 overexpression, as demonstrated in our study.

Importantly, our results not only reveal structural synaptic loss upon BIN1iso1 expression but also demonstrate functional consequences on synaptic transmission. In *Drosophila*, thanks to ERGs, we showed an early progressive loss of synaptic transmission between photoreceptor neurons and the optic lobe. In rat primary neuronal cultures, we used MEA technology to assess neuronal activity. A key advantage of using this technology is its ability to measure neuronal activity directly, without relying on indirect calcium or glutamate sensors for example. For all these recordings, changes in signal frequency and amplitude can result from alterations in either intrinsic neuronal excitability or synaptic transmission, making it challenging to distinguish between the two. One way to disentangle these effects is by analyzing network synchronicity, which has been used as an indirect measure of synaptic transmission. This way, BIN1 knockout iPSC-derived neurons were shown to be defective in functional synaptic connectivity (Saha et al., 2024). To go further, in this work, we combined MEA with microfluidic compartmentalization to directly assess functional connectivity within a defined network of pre- and postsynaptic neuron populations. While we did not observe changes in firing rate or spike amplitude in the presynaptic chamber where BIN1iso1 was overexpressed, we did detect a clear reduction in functional connectivity as the pre-to-post synaptic drive was close to 1 upon BIN1iso1 expression. We did not observe the same result with BIN1iso9, reinforcing the isoform-specific effect. The absence of effect on the firing rate or spike amplitude contrasts with previous reports showing hyperexcitability with increased spike frequency upon BIN1 overexpression (Voskobiynyk et al., 2020). This may be explained by differences in the maturity of the neurons at the time of transduction and analysis. More broadly, our findings further support BIN1’s role as a modulator of synaptic transmission and network dynamics.

Our results consistently point to a presynaptic-specific role for BIN1iso1. BIN1 has been shown to be predominantly localized in the pre- and post-synaptic compartments in rat primary neuronal cells and mouse brain (De Rossi et al., 2020; Schürmann et al., 2020). In both the *Drosophila* photoreceptor model and the larval NMJ model, BIN1iso1 was tested presynaptically. We further confirmed this compartment-specific effect using a microfluidic device allowing for precise control of protein expression in rat primary neuron cultures. This system revealed a clear reduction in synaptic density when BIN1iso1 was expressed presynaptically, but not postsynaptically. This is in accordance with a role of BIN1 in regulating presynaptic synaptic vesicle release probability seen in conditional knockout mouse brain (De Rossi et al., 2020), and with a presynaptic role of BIN1 in the synaptic vesicle endocytic cycle in inhibitory synapses of mouse primary neuronal cultures (Barata et al., 2025). A postsynaptic role has also been reported in the exocytosis of GluA1 AMPA receptor subunit (Schürmann et al., 2020). These observations suggest that BIN1 may have distinct, context-dependent functions on both sides of the synapse.

Mechanistically, the synaptic toxicity of BIN1iso1 appears to be linked to disruptions in endo-lysosomal trafficking. Indeed, BIN1iso1 expression led to the accumulation of endosome-like large vesicular structures in the terminals of photoreceptor neurons, as observed via immunofluorescence and electron microscopy. In addition, the synaptic overgrowth phenotypes, especially the formation of satellite boutons at the larval NMJ, have been reported in mutants affecting endocytosis, such as *endophilin, synaptojanin, dynamin, AP180, synaptotagmin* and *σ2-adaptin* (Choudhury et al., 2016; Dickman et al., 2006). These morphological changes are thought to arise from defective intracellular trafficking of signaling molecules from the BMP pathway that regulate synaptic growth (O’Connor-Giles et al., 2008). In addition, BIN1iso1-induced photoreceptor degeneration was previously linked to trafficking defects and could be suppressed by modulating Rab5 or Rab11 activity (Lambert et al., 2022), both key regulators of early and recycling endosomes. In the present study, overexpression of Rab11 only partially rescued in an additive manner the synaptic overgrowth phenotype induced by BIN1iso1, suggesting mechanistic divergence between the soma and presynaptic terminals. Similarly, Rab11 overexpression has been shown to mitigate synaptic toxicity in *Drosophila* models of Parkinson’s disease involving *Parkin* and *PINK1* (Rai et al., 2023), yet it does not rescue overgrowth phenotypes in *AP-2* loss-of-function *Drosophila* models (Choudhury et al., 2022). The existence of different mechanisms in subcellular compartments in mammals may also explain why BIN1 loss-of-function has been associated with both smaller and bigger Rab5-positive endosomes in rodent primary neurons (Barata et al., 2025; Calafate et al., 2016). Of note, we also observed that Rab11 overexpression alone reduced bouton number at the *Drosophila* NMJ, a novel finding consistent with previously reported increases in bouton number in Rab11 loss-of-function mutants (Khodosh et al., 2006). The endo-lysosomal dysregulation by BIN1 is of interest for AD as endo-lysosomal defects are considered potentially causal in AD (Nixon et al 2024). BIN1 is known to interact with many components of this system, such as RIN3, another AD genetic risk factor (Dourlen et al., 2025; Kajiho et al., 2003; Shen et al., 2020). Together, these data further support a role for BIN1iso1 in regulating synaptic architecture via intracellular endo-lysosomal trafficking pathways.

Altogether, our data establish BIN1iso1 as a potent modifier of synaptic integrity, at structural and functional levels. The early synaptic alterations, the accumulation of endosomal vesicles, and the isoform and compartment specificity of BIN1iso1 toxicity provide novel insights into the molecular mechanisms by which BIN1 may contribute to neurodegeneration. These findings are particularly relevant given the genetic association of BIN1 with Alzheimer’s disease and suggest that isoform-specific targeting could represent a therapeutic strategy.

## Acknowledgements

The authors would like to acknowledge the BioImaging Center Lille (BICeL) and la PLateforme d’Expérimentation et de Haute Technologie Animale (PLEHTA), both part of Plateformes Lilloises en Biologie et Santé (PLBS) - UAR 2014 - US 41), for microscopy maintenance and animal housing, respectively. pLenti.PGK.LifeAct-Ruby.W and pLenti.PGK.LifeAct-GFP.W were gifts from Rusty Lansford (Addgene plasmid # 51009; http://n2t.net/addgene:51009; RRID:Addgene_51009 and Addgene plasmid # 51010; http://n2t.net/addgene:51010; RRID:Addgene_51010). The authors thank the vectorology service of the University of Bordeaux for lentiviral packaging. The 4F3 and nc82 antibodies developed by Goodman, C. (University of California, Berkeley) and Buchner, E. (Universitaetsklinikim Wuerzburg) were obtained from the Developmental Studies Hybridoma Bank, created by the NICHD of the NIH and maintained at The University of Iowa, Department of Biology, Iowa City, IA 52242. Stocks obtained from the Bloomington *Drosophila* Stock Center (NIH P40OD018537) were used in this study.

This work was supported by the Joint Programme ̶ Neurodegenerative Disease Research (JPND; 3DMiniBrain), by France Alzheimer (#328 ADhesion, #1999 BIN1DROSO), by Fondation Vaincre Alzheimer (LECMA Grant The BIN1-Tau neurotoxic link in *Drosophila*), by ANR (TAUFUNALZ project, ANR-23-CE14-0064), by the LabEx (Laboratory of Excellence) DISTALZ (Development of Innovative Strategies for a Transdisciplinary approach to Alzheimer’s disease ANR-11-LABX-01), by the Région Hauts-de-France Regional Council and by an Alzheimer’s Association Grant (AARG-22-926152). CG was supported by a fellowship from FranceAlzheimer (Soutien aux jeunes chercheurs – AAP JC 2024 – #6473). This work has been partially undertaken with the support of IEMN CMNF facilities and supported by the French Renatech network.

## Availability of data and material

Data and material from the current study are available from the corresponding authors upon request.

