## supplementary materials for "BIN1 expression in the presynaptic compartment leads to isoform-specific synaptotoxicity"

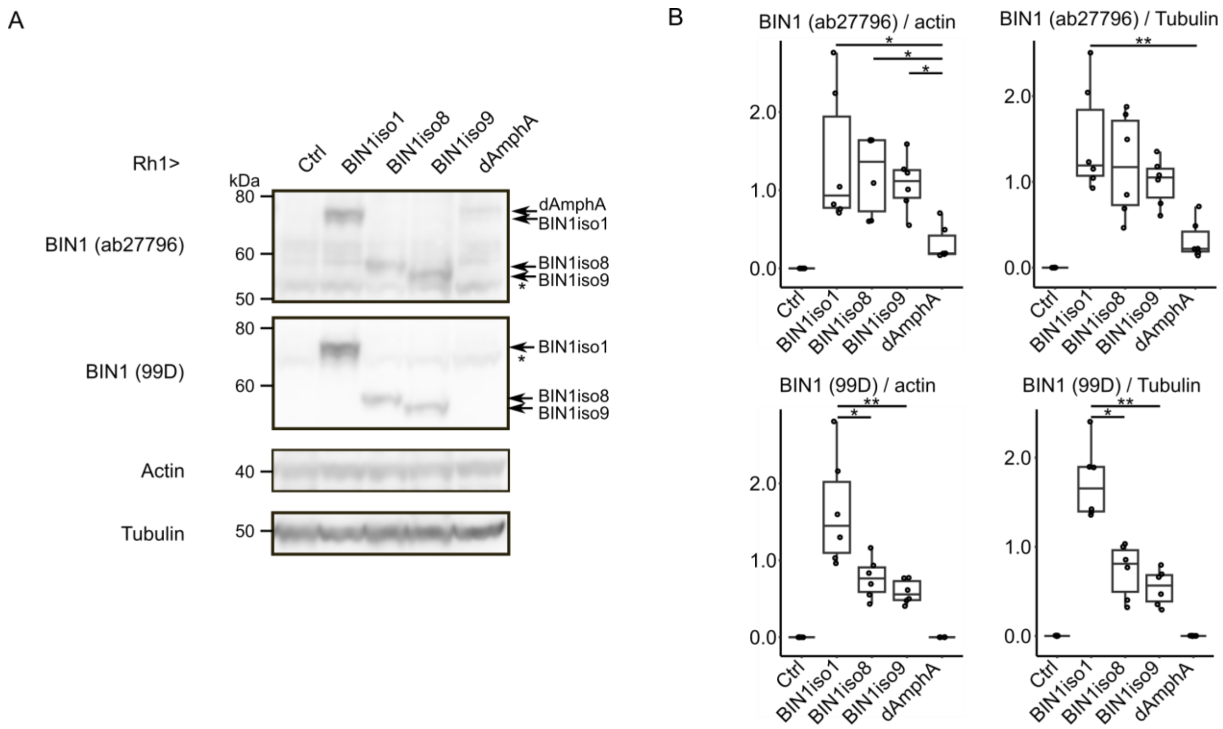


**Supplementary Figure 1:** Analysis of BIN1 levels in *Drosophila* expressing human BIN1 isoforms and *Drosophila* Amphiphysin isoformA under a Rh1 driver. **A.** Representative Western blots analyzing the transgenic expression of human BIN1 isoforms and *Drosophila* Amphiphysin in 1-2 day-old *Drosophila* heads. BIN1 was detected using two antibodies (ab27796 and 99D). Actin and Tubulin were used as loading controls. Non-specific bands were marked with stars. **B.** Quantification of data obtained from 6 independent experiments. Statistical analyses were performed with non-parametric Kruskal Wallis tests followed by Dunn's multiple comparisons (*p*-values adjusted using the Holm method, * $p<0.05$; ** $p<0.01$). The Ctrl group and the Ctrl and dAmphA groups were excluded from the statistical analysis for blots obtained using the BIN1 ab27796 and BIN1 99D antibodies, respectively.

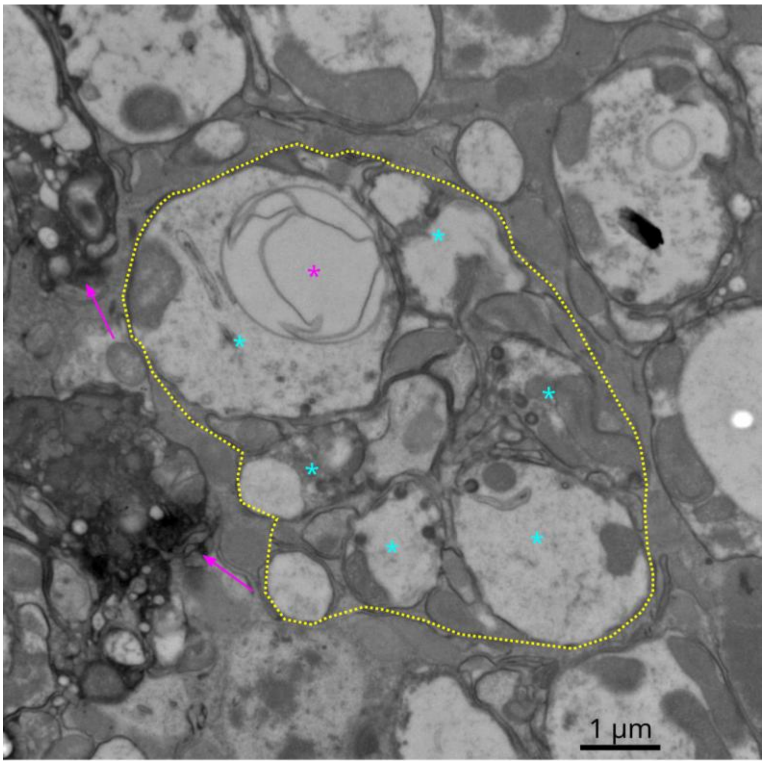


**Supplementary Figure 2:** Electron microscopy image showing the degeneration of synaptic terminals in *Drosophila* photoreceptor neurons expressing BIN1 iso1 at day 15. Six synaptic terminals (cyan stars) are gathered into a lamina cartridge (yellow dotted line). Expression of BIN1 iso1 resulted in the presence of giant vesicles (magenta star). Magenta arrows point to degenerating synaptic terminals in neighboring cartridges.

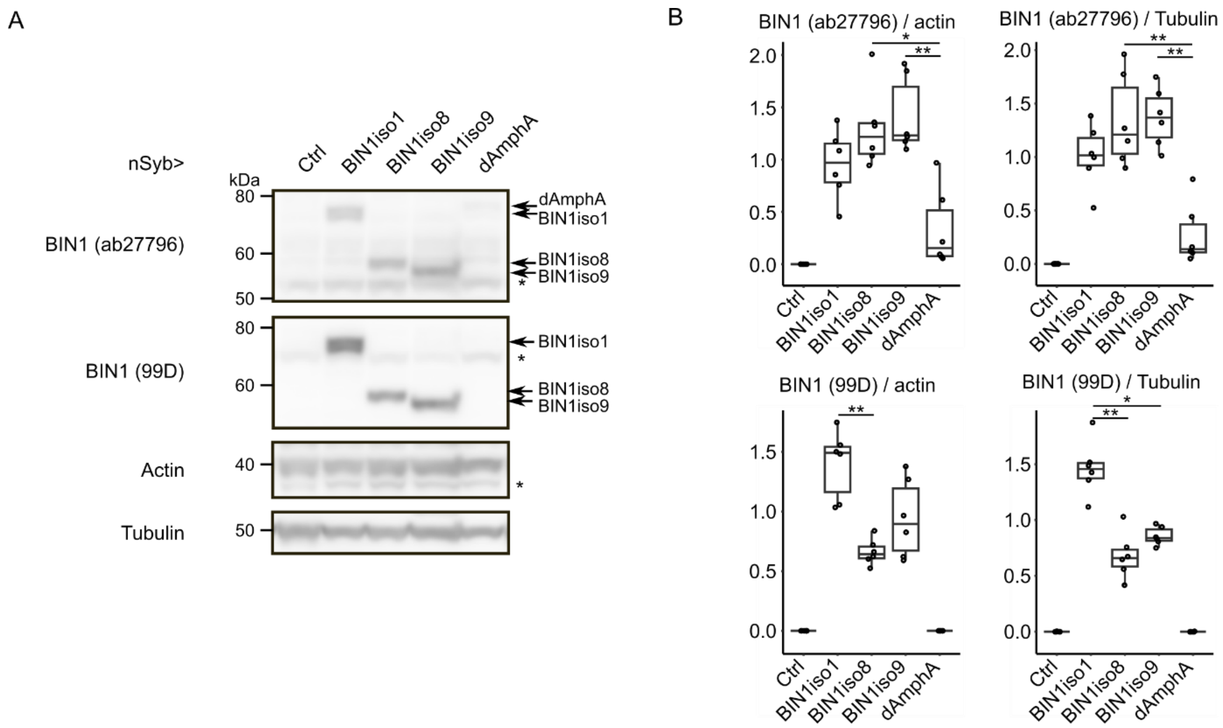


**Supplementary Figure 3:** Analysis of BIN1 levels in *Drosophila* expressing human BIN1 isoforms and *Drosophila* Amphiphysin under a nSyb driver. **A**. Representative Western blots analyzing the transgenic expression of human BIN1 isoforms and *Drosophila* Amphiphysin in 1-2 day-old *Drosophila* heads. BIN1 was detected using two antibodies (ab27796 and 99D). Actin and Tubulin were used as loading controls. Non-specific bands were marked with stars. **B.** Quantification of data obtained from 6 independent experiments. Statistical analyses were performed with non-parametric Kruskal Wallis tests followed by Dunn's multiple comparisons (*p*-values adjusted using the holm method, * p<0.05; ** p<0.01). The Ctrl group and the Ctrl and dAmphA groups were excluded from the statistical analysis for blots obtained using the BIN1 ab27796 and BIN1 99D antibodies, respectively.

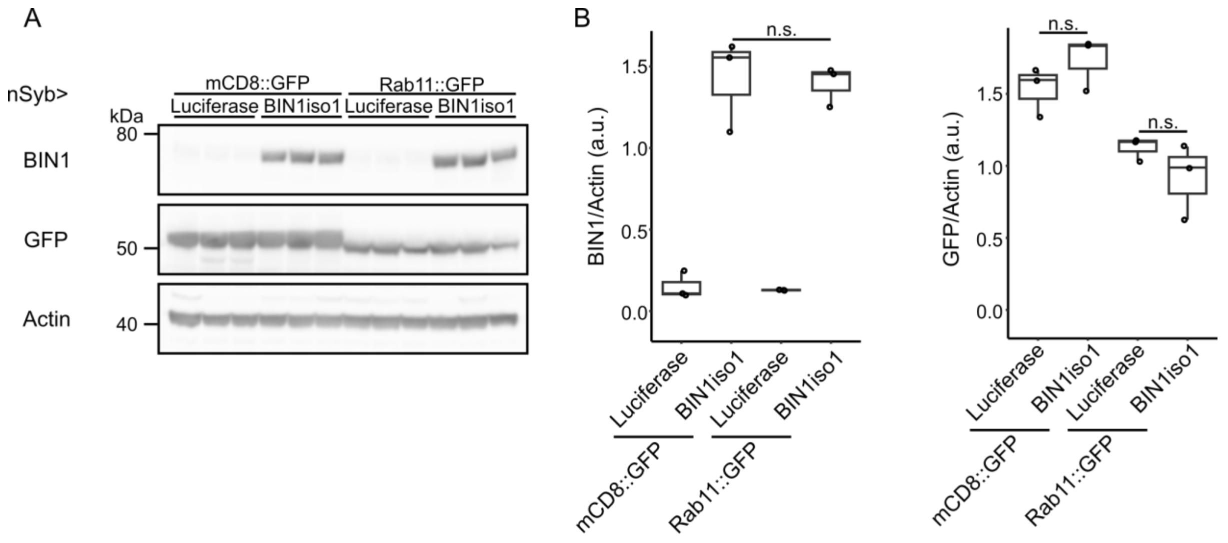


**Supplementary Figure 4:** Protein levels in *Drosophila* expressing BIN1iso1 and Rab11. **A.** Representative Western blots using *Drosophila* head protein extracts from animals expressing luciferase, mCD8::GFP (both used as controls), BIN1iso1 and Rab11::GFP, under a Nsyb driver, using anti-BIN1 (99D), anti-GFP and anti-actin antibodies. **B.** Quantification of BIN1 and GFP from 3 independent experiments. Statistical analyses were performed with non-parametric Kruskal Wallis tests followed by Dunn's multiple comparisons (*p*-values adjusted using the holm method, n.s. not significant).

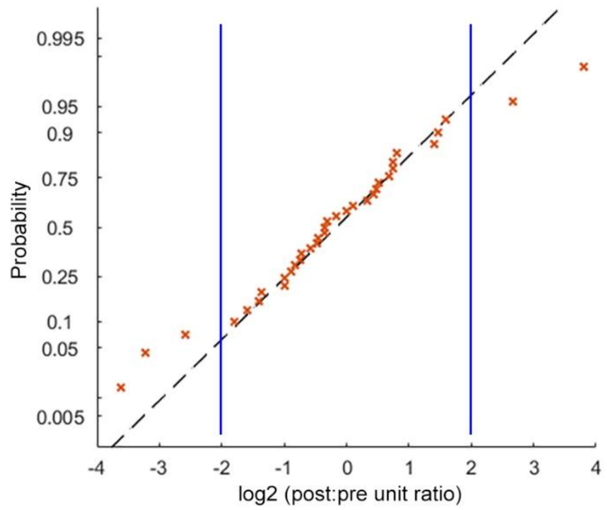


**Supplementary Figure 5:** Probability plot comparing the distribution of the fold differences between unit numbers detected in presynaptic and postsynaptic chambers to the normal distribution. Each point indicates one MEA-integrated microfluidic device, in which the spike sorting algorithm detected at least one unit in both chambers (n = 35 devices from 7 independent cultures). Dashed line passes through the lower and upper quartiles. Solid lines indicate the limits for 4-fold difference in unit numbers and correspond to the exclusion criteria used for this dataset (|*z*-score| = 1.5).

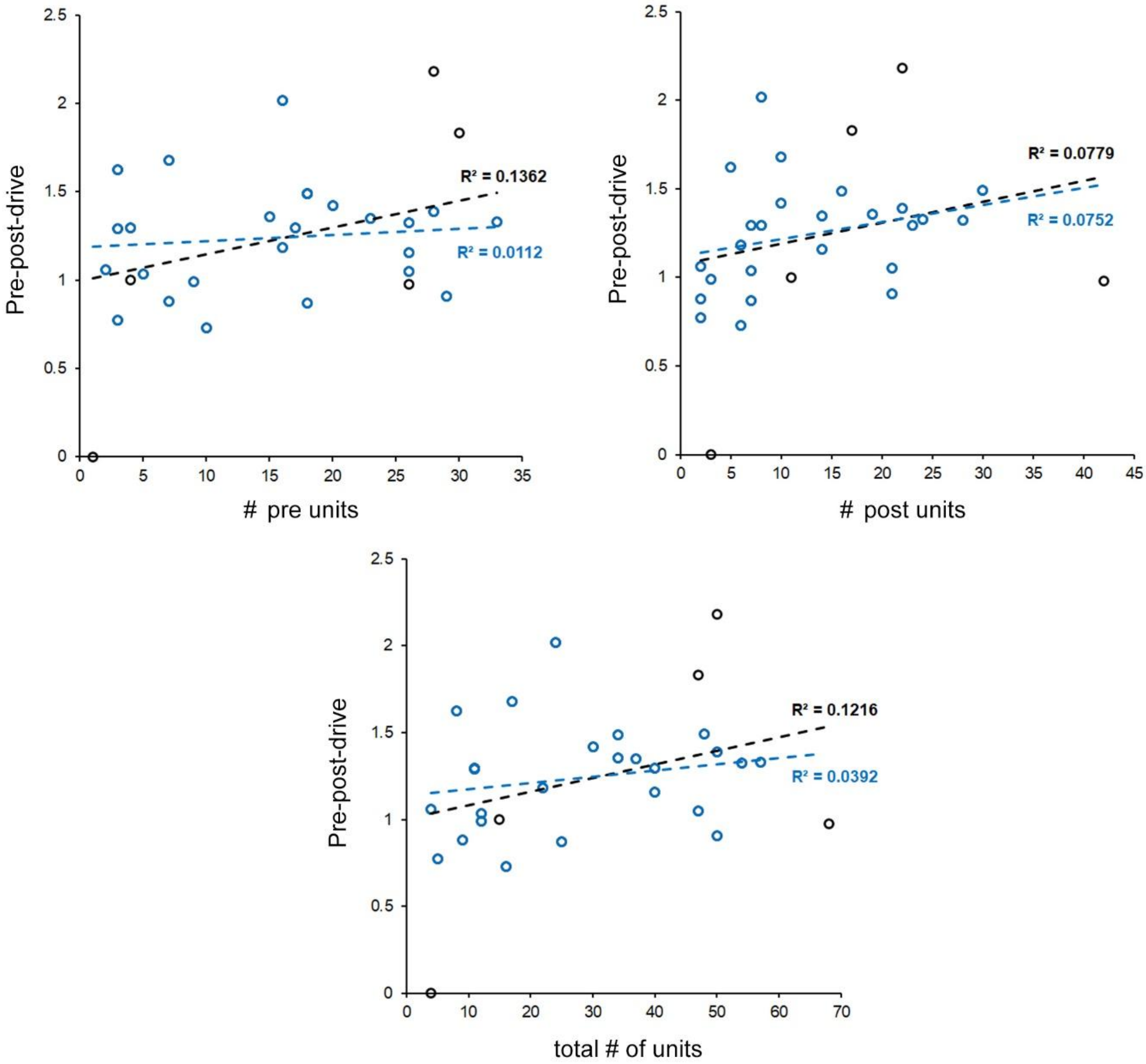


**Supplementary Figure 6:** Correlation between detected unit numbers and *pre-post-drive*. Each point indicates one MEA-integrated microfluidic device within the normal distribution of fold differences in unit numbers (n = 30 devices from 7 independent cultures). Outlier data points (black dots) were included for clarity. Dashed lines show the least-squares fits before (black) and after (blue) outlier exclusion.
